# Inference of CRISPR Edits from Sanger Trace Data

**DOI:** 10.1101/251082

**Authors:** Tim Hsiau, David Conant, Nicholas Rossi, Travis Maures, Kelsey Waite, Joyce Yang, Sahil Joshi, Reed Kelso, Kevin Holden, Brittany L Enzmann, Rich Stoner

## Abstract

Efficient precision genome editing requires a quick, quantitative, and inexpensive assay of editing outcomes. Here we present ICE (Inference of <u>C</u>RISPR <u>E</u>dits), which enables robust analysis of CRISPR edits using Sanger data. ICE proposes potential outcomes for editing with guide RNAs (gRNAs) and then determines which are supported by the data via regression. Additionally, we develop a score called ICE-D (Discordance) that can provide information on large or unexpected edits. We empirically confirm through over 1,800 edits that the ICE algorithm is robust, reproducible, and can analyze CRISPR experiments within days after transfection. We also confirm that ICE strongly correlates with next-generation sequencing of amplicons (Amp-Seq). The ICE tool is free to use and offers several improvements over current analysis tools. For instance, ICE can analyze individual experiments as well as multiple experiments simultaneously (batch analysis). ICE can also detect a wider variety of outcomes, including multi-guide edits (multiple gRNAs per target) and edits resulting from homology-directed repair (HDR), such as knock-ins and base edits. ICE is a reliable analysis tool that can significantly expedite CRISPR editing workflows. It is available online at ice.synthego.com, and the source code is at github.com/synthego-open/ice

## Introduction

CRISPR is a precise programmable tool used for genome editing. Because CRISPR technology is relatively easy to use compared to other genome editing tools, it has increased in popularity in recent years. CRISPR involves a guide RNA (gRNA) that binds to genomic locus and a Cas nuclease that creates a double-stranded break (DSB) at the targeted site. After a DSB is made, non-homologous end joining (NHEJ) can introduce insertions or deletions (indels) that potentially knock out the targeted gene. Alternatively, if a DNA donor template is provided, knocking in a sequence of interest is possible through homology-directed repair (HDR). The editing outcomes of a CRISPR experiment are unpredictable and result in a heterogeneous cell population. Additionally, it is not readily apparent to the researcher if an edit has occurred and whether or not to continue culturing the edited cells. Various methods have been developed to address the problem of identifying the sequences present in an edited population at the targeted locus.

Previous methods to infer the composition of an unknown mixture of sequences have included next-generation sequencing (NGS) of amplicons (Amp-Seq), Tracking Indels by Decomposition (TIDE) [1], and compressed sensing [2]. Amp-Seq offers sequence-level resolution and more sensitivity, but it is less widely available, has a longer turnaround time, and comes at a higher cost per sample unless a substantial number of samples are batched.

A significant benefit of the TIDE approach is that only Sanger sequence data are required for analyzing mixed populations, avoiding the need for NGS. While TIDE has been successful in reducing the barriers to entry for CRISPR, several characteristics of the program hinder its ease of use and automatability. For instance, users have to manually tune algorithm parameters and process each sample (a batch processing version of the algorithm is not readily available). Both of these aspects make TIDE a time-consuming analysis. Additionally, TIDE is limited to assessing editing outcomes in which an individual guide is used and cannot handle experiments where multiple guides are used simultaneously.

To develop a fast and robust method for verifying CRISPR edits, we focused on improving the procedure set forth in TIDE. In the TIDE method, the targeted locus is amplified and Sanger-sequenced for both control and edited samples. Ideally, the edit site is within 200-300 bp of the sequencing primer, resulting in a read where the first hundred base pairs are of high quality and the following bases are potentially mixed (indicating an edited population with heterogeneous outcomes). Computationally, different editing proposals are generated using the control sample, and regression is used to compute how much of each editing outcome is observed in the mixed Sanger read.

Here we present an improved algorithm, called ICE (Inference of CRISPR Edits), that addresses the drawbacks of TIDE and makes the editing analysis process more robust and automated. The tool can be used to assess experiments in which one guide or multiple guides are used to edit a targeted locus and is capable of analyzing edits resulting from NHEJ (i.e., knockouts) as well as HDR (knock-ins, base edits).

## Methods

### ICE Algorithm

We re-implemented the TIDE [1] algorithm in Python and then added improvements to support more analysis cases and to make the algorithm more robust. Figure 1 shows the algorithm flowchart that corresponds to the steps below.

**Figure 1.**
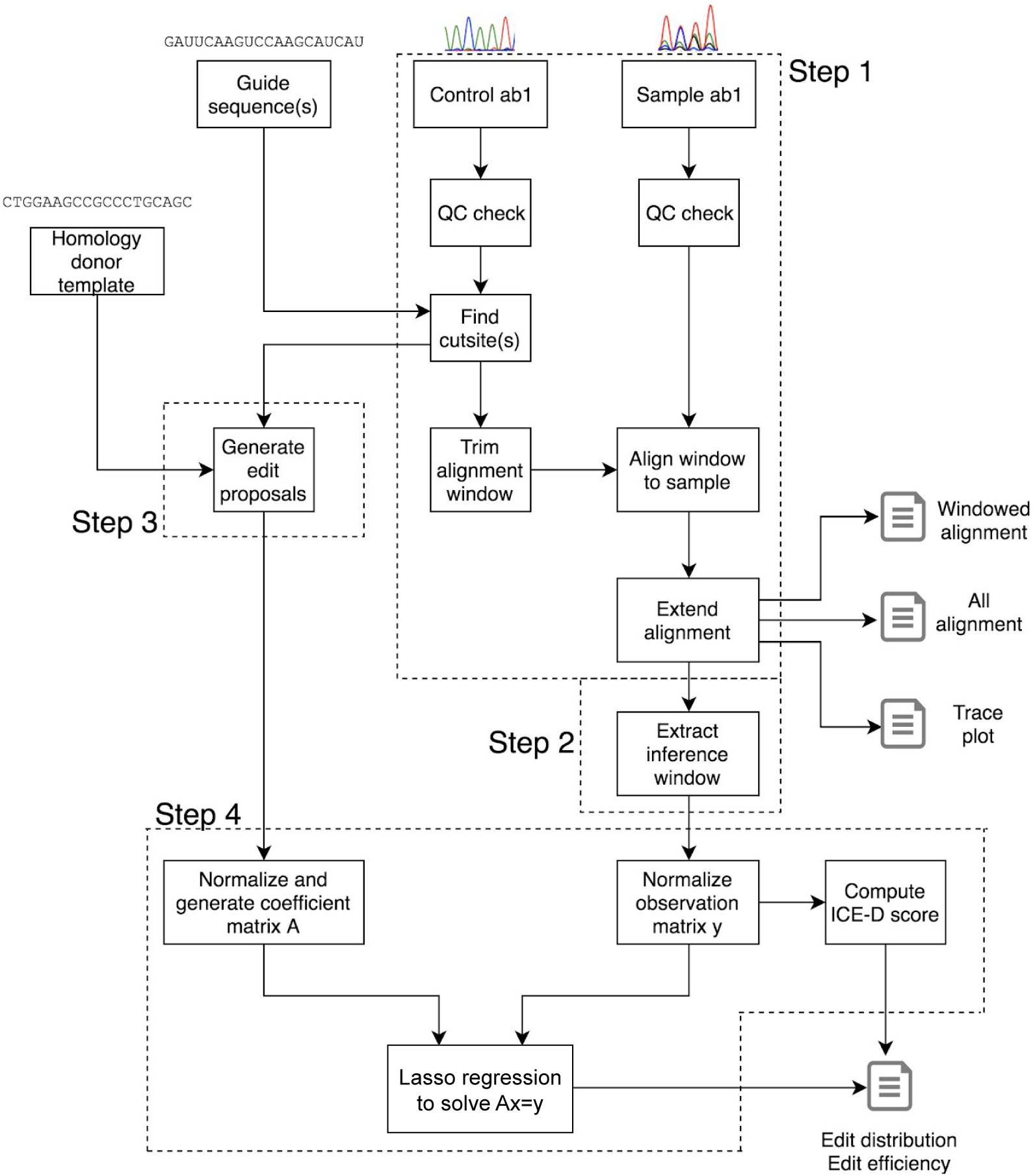
Algorithm flowchart. The inputs to the algorithm (top) are the control ab1 file, the sample ab1 file, the guide sequence(s), and an optional DNA donor or base editor template. The algorithm checks the data quality, generates edit proposals, and then runs a Lasso regression to identify which edit proposal sequences are most likely present in the sample. The program then outputs the quality of the results, the percentage of the sample population that has been edited, and the identity of the edited sequences.

Step 1: We align the two trace files by finding a high-quality window of the control trace upstream of the cut site and trimming it to end 15 bp upstream of the cut site. This alignment window is defined as a region of the Sanger trace that has a windowed average with Phred quality scores of greater than 30. The alignment window in the control is then aligned against the edited sample (Fig 2). By ignoring the poor quality bases often found at the very beginning of a Sanger trace, we found that this alignment method is robust and scales well for reliably processing many ab1 files.

**Figure 2.**
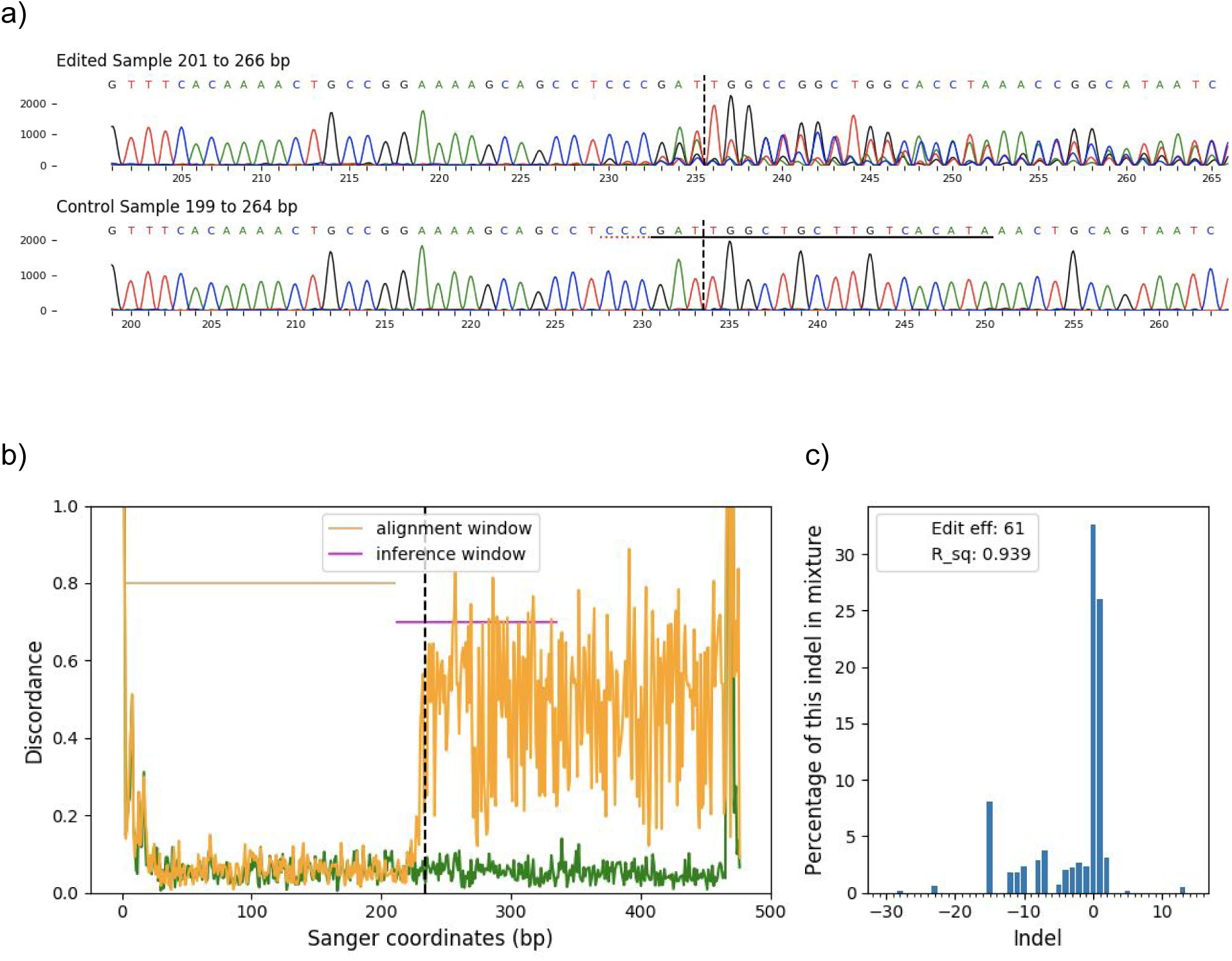
An example of the outputs from the ICE software for a guide targeting the human gene *GRK5*. **a)** Trace file segments spanning the cut site from the control and the edited samples are generated for every analysis. The guide sequence is underlined in black, and the PAM sequence is denoted by a dotted red underline in the control sample. Vertical dotted lines denote the expected cut site. **b)** Discordance for the edited (orange) and control (green) trace files. The vertical dotted line marks the cut site. The alignment window marks the region of the traces with high Phred scores and is used to align the edited and control traces. The inference window marks the region of the traces around the cut site which will be used to infer the change in sequence between the edited and control traces. Visualizing this way, we can see that discordance is a robust signal of trace irregularity that can approximate the location and prevalence of editing. c) Indels and their relative prevalence for this example, as calculated by ICE.

Step 2: We defined the inference window to be the segment of the trace data used for the regression. The inference window starts 25 bp upstream of the cut site and extends up to 100 bp downstream of the cut site. The algorithm trims the inference window based on the quality score of the control sample. We limit the inference window length, as adding extra bases has diminishing returns and can hurt the regression as Sanger sequencing quality decreases over the length of the read.

Step 3: We also made improvements to the edit proposal process, a step in which the algorithm generates potential post-editing genome sequences for use in the analysis. We aimed to support the analysis of many use cases for precision genome engineering, including the following:

1. <u>Editing with one guide RNA:</u> the algorithm uses a default indel range of deletions of up to 30 bp and insertions of up to 14 bp to generate a list of potential edits (sequences and traces). For deletions, the associated trace data for that base is deleted, while for insertions a uniform distribution of 25% for each base is inserted. The trace data for other bases are copied from the wild-type. These simulated traces are then used in the Lasso regression (step 4 below).
2. <u>Editing a single locus with multiple guide RNAs simultaneously (multi-guide):</u> the algorithm first generates edit proposals for each guide as above; then, for every pair of guides, the algorithm generates edit proposals which cover the following two classes: a) both guides cut independently with indel formation or b) both guides cut “in tandem” and the intervening sequence is omitted.
3. <u>Knock-ins/ Homology-directed repair:</u> the algorithm aligns the input template with the control sequence and then for any bases that differ in the alignment are simulated by a score of 97% for the desired base and 1% for each of the three other bases. The alignment expects at least 15 bp of sequence on both ends of the template to match the genomic target.
4. <u>Base editing:</u> the same method as in the homology-directed repair case can be used to analyze samples. It would be necessary to input a template sequence that mimics the expected base editing outcomes.

Step 4: After the edit proposal stage, a regression is performed to infer the frequencies of each proposal sequence. In the regression, x is solved for in the equation Ax=y, where A is a matrix composed of the simulated traces and y is the observed outcomes vector (the edited sequencing trace). Lasso regression finds a linear combination of the edit proposals that best explains the observed trace of the edited sample. Lasso regression mitigates overfitting to the noise in sanger data via L1 regularization, producing more parsimonious results compared to alternative regression algorithms like non-negative least squares. The relative prevalence of each edit proposal is extracted from the vector of weights of the regression (x). Percentages of individual edits are rounded to the nearest whole percentage point to reflect the model’s underlying confidence about the accuracy of contribution estimations. The correlation between the regression derived and the observed edited sequencing trace (r^2^) measures the extent to which the edit proposals can explain the edited sequencing trace.

ICE outputs several graphs and metrics that indicate the quality of the results and the frequency of different editing outcomes (Fig 2).

### ICE-D Algorithm

We next wanted to account for large insertions or deletions that are not expected and would not be accounted for by any edit proposal. We noticed that when samples edited by multiple guides were analyzed with only one guide, there was a low ICE r^2^ value and low reported editing. Upon manual inspection, however, we found that a large proportion of the edited Sanger trace was discordant with the control.

To account for unexpected edits, we created a new program that assesses the average discordant signal, called ICE-D (ICE-Discordance). ICE-D is motivated by the observation that given a high-quality alignment upstream of the cut site, we could assume that any signal downstream of the cut site discordant with the control trace should be evidence of editing. The ICE-D score is proportional to the discordance score in the inference window (Fig 2). We calibrated the ICE-D correction factor by finding the indel percentage and the average discordant signal for 1805 edits and then performing a least squares linear regression between the two metrics.

### Program Outputs

For interpreting results and checking algorithm settings, ICE outputs summary JSON and xlsx spreadsheets, plots, and alignments.

### Source Code

The source code is available at github.com/synthego-open/ice, a docker container is on the docker hub at synthego/ice, and a publicly accessible webtool can be found at http://ice.synthego.com.

### Editing of Cell Cultures

Editing was performed with Synthego synthetic single guide RNAs (sgRNAs) on a variety of cell lines. In general, sgRNAs were synthesized with or without modifications and complexed with Cas9 at a molar ratio of 9:1 (sgRNA:Cas9) to form ribonucleoproteins (RNPs). The resulting RNPs were transfected into the respective cell line using a Nucleofector^TM^ (Lonza; Basel, Switzerland). Transfected cells were recovered in normal growth medium, plated into 96-well plates, and incubated in humidified 37°C/5% CO_2_. After 48 hours, cells were lysed and genomic DNA was extracted from the cells using QuickExtract™ DNA Extraction Solution (Lucigen; Middleton, WI) to each well of the plate. Knock-in editing was performed in HEK293 cells using modified sgRNAs (Synthego) and single-stranded DNA (ssDNA) donor templates (Eurofins Genomics). The ssDNA templates were designed to knock in sequences of varying length (+0 (SNP), +14 or +36 bp) with symmetric 40 bp homology arms. The components were introduced at a ratio of 9:1:3 (sgRNA:Cas9:ssDNA). All other experimental details were the same as the non-HDR experiments.

### Sanger & Next-Generation Sequencing

We targeted 32 genes each with three sgRNAs specifically designed to produce one or more large deletions (multi-guide). For each gene, the guides were transfected individually and together, for a total of 128 samples. Three to four replicate edits were performed and Sanger-sequenced. One replicate was amplified for Next-Generation Sequencing of Amplicons (Amp-Seq).

For Sanger sequencing samples, PCR primers were designed to amplify a 500-800 bp segment containing the cut site. PCR was performed on lysed genomic samples using Taq polymerase. Sequencing was then performed through a commercial vendor (Sequetech; Mountain View, CA) with one of the two primers used for amplification. For HDR transfections, we designed primers such that the same set could be used for both Sanger and Amp-Seq. The resulting amplicons were between 300-500 bp with the cut site 100 bp from the forward primer. One gene, comprising four samples, repeatedly failed Sanger sequencing and was dropped from subsequent analysis.

For Amp-Seq samples, a 200-300 bp segment containing the cut site was PCR-amplified. The amplicons were purified, quantified using Nanodrop^TM^, and sent to the MGH DNA core facility for their CRISPR sequencing service. A summarization analysis was performed using the MGH NGS data pipeline, which reported the sequence and abundance of amplicons.

The ICE replicates were averaged together to compare with the Amp-Seq findings. Indels were summarized by length and then compared. ICE-predicted sequences were also compared with contigs assembled from Amp-Seq data.

### Comparison of ICE and TIDE

Three hundred and forty-two samples with high r^2^ values (>0.95) were chosen from 1805 samples that were analyzed using ICE. All edits were conducted using a single guide and samples were chosen to span a range of indel percentages. These samples were then manually analyzed using the TIDE website [3] in August 2018. Default parameters for the TIDE website were first attempted, and subsequent parameter tuning was performed if the initial setup resulted in a failed analysis. The overall indel percentage (editing efficiency) was then compared between TIDE and ICE.

### Simulated Homology-Directed Repair and Base Editing

The SNP rs2072579 was amplified from PGP1 (George Church’s induced pluripotent stem cell line) and HEK293 genomic lysate. The amplicons were first sequenced to verify that the PGP1 cell line was homozygous G/G and HEK293 cell line was homozygous C/C. Amplicon masses were quantified using a Fragment Analyzer (AATI; Ankeny, Iowa) and then hand-pipetted to generate mixes to simulate different editing outcomes ranging from 5% to 95% single base editing.

## Results

The ICE software is easy to use with no parameter tweaking required. The default indel limits are set at −30 and +14 for a single DNA break, which should cover most use cases. The software also outputs files that aid the user in checking the quality of edits and the interpretation of results (Fig 2).

We confirmed that our ICE tool was robust by performing an analysis on a batch of 1805 edits performed over multiple experiments. Our ICE tool takes on average four seconds to process each sample on a laptop in single threaded mode (MacBook Pro 2017). We also used this batch of edits to calibrate our ICE-D correction factor. The ICE-D correction factor is multiplied by the average discordant signal downstream of the cut site in the inference window to yield a proposal-agnostic guess of the indel percentage present in the cell.

We then tested the ability of ICE-D to detect unexpected edits by re-running the analysis and only supplying one guide sequence for multi-guide samples. When only one guide is provided, the edit proposal stage does not generate all edit outcomes possible for a multi-guide experiment. The lack of all possible edit outcomes then becomes an issue if those sequences are actually present and contribute to the Sanger signal. In those cases, both ICE and TIDE will give a low goodness-of-fit metric and may underestimate the actual indel percentage. However, ICE-D is still able to detect editing as can be seen by the gray dots (signifying r^2^ ≤ 0.95) of Figure 3b.

**Figure 3.**
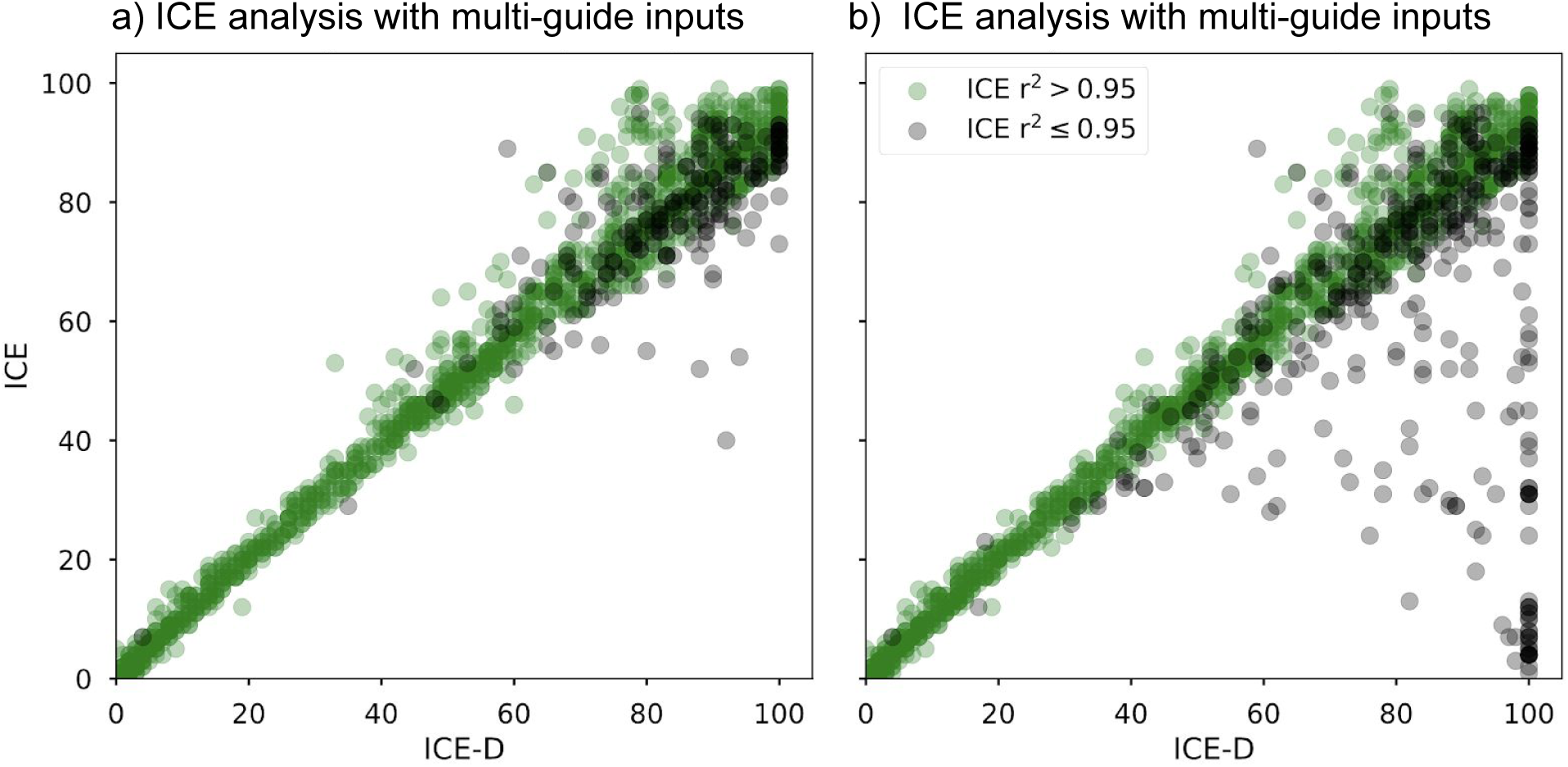
The ICE-D empirical correction factor was calibrated using ICE results from 1805 edits. **a)** ICE analysis performed with multi-guide sequence inputs. **b)** ICE analysis without multi-guide inputs provided. For the analysis with multi-guide information, 78% of the samples have high (>0.95) ICE r^2^, and that number drops to 71% for the analysis without the multi-guide information. When multi-guide information is not provided, ICE-D is able to detect unexpected edits when the ICE goodness of fit is low (<0.95) and ICE-D score is high (gray data points in the bottom right of graph b).

To validate our algorithm, we compared ICE results with results from TIDE and next-generation sequencing of amplicons (Amp-Seq). First, we analyzed 342 samples through both TIDE and ICE using default settings. These samples were chosen to span a range of 0-95% editing efficiencies and for having a high r^2^ in ICE. Figure 4 shows that the correlation between ICE and TIDE results is high with an r^2^ of 0.99 (A). The correlation between ICE-D and TIDE is slightly lower (r^2^ =0.97; B). Although TIDE and ICE may output similar results, TIDE often requires users to tune the parameters for alignment to be able to get an interpretable result. ICE is fully automated and does not require users to manipulate any parameters.

**Figure 4.**
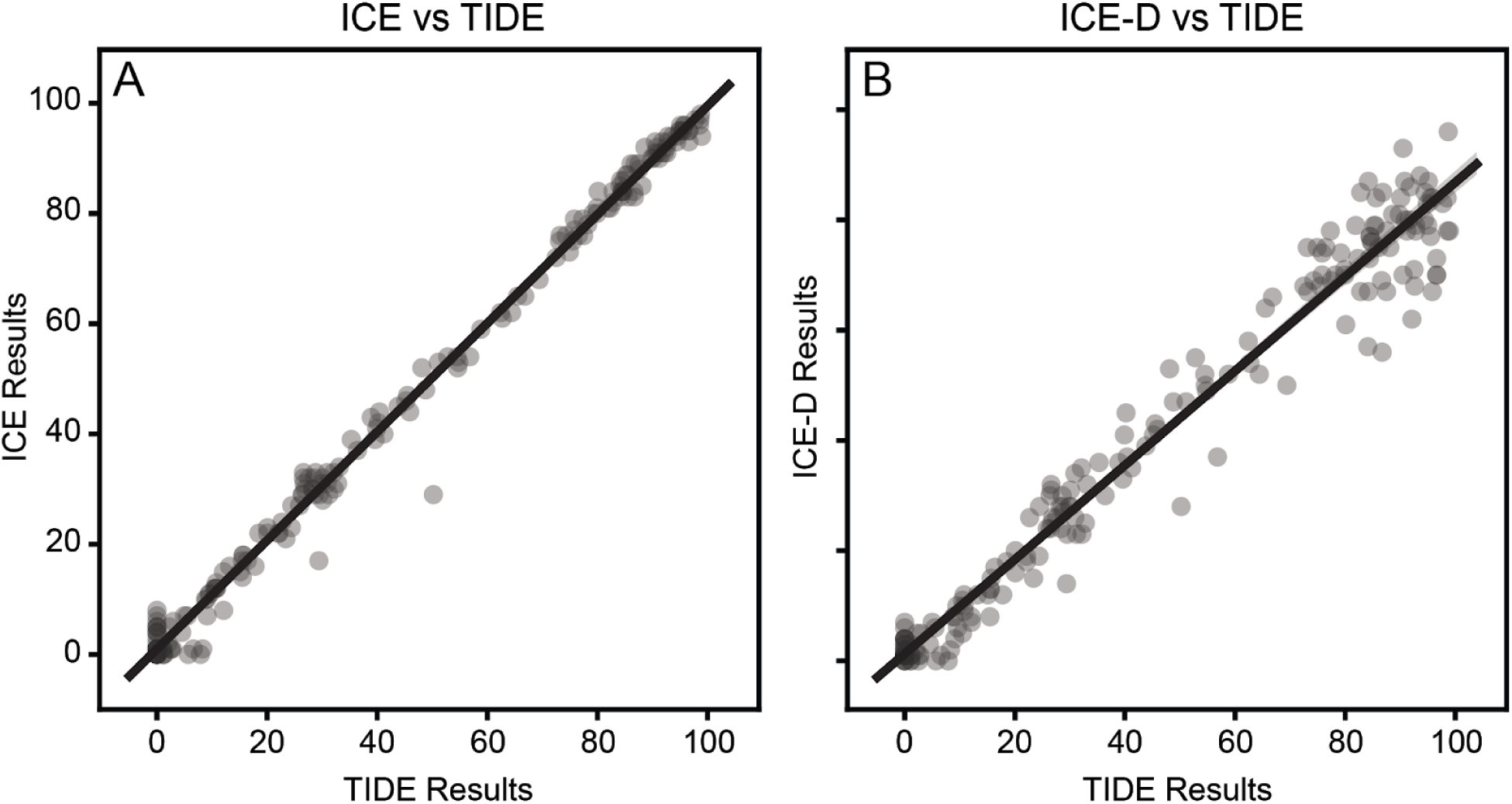
**A)** ICE results strongly correlate with TIDE results (r^2^= 0.99). **B)** ICE-D relies on more assumptions and has a slightly poorer correlation with TIDE, r^2^= 0.97.

We next sought to show that ICE correlates well with Amp-Seq, the current gold standard in CRISPR genomic analysis. We performed amplicon sequencing on 92 samples using MGH CRISPR sequencing service. We correlated the ICE results with the Amp-Seq results for each indel size in all samples. Individual examples of this correlation are shown in Figure 5 A-D. We found a high correlation, with an overall r^2^ = 0.93 (Fig 5 E-F). To show that the ICE results are correct at the sequence level, instead of a summarized indel size, we also manually inspected the sequences from Amp-Seq and ICE. In Appendix A, we have some examples of sequences predicted by ICE matching Amp-Seq results.

**Figure 5.**
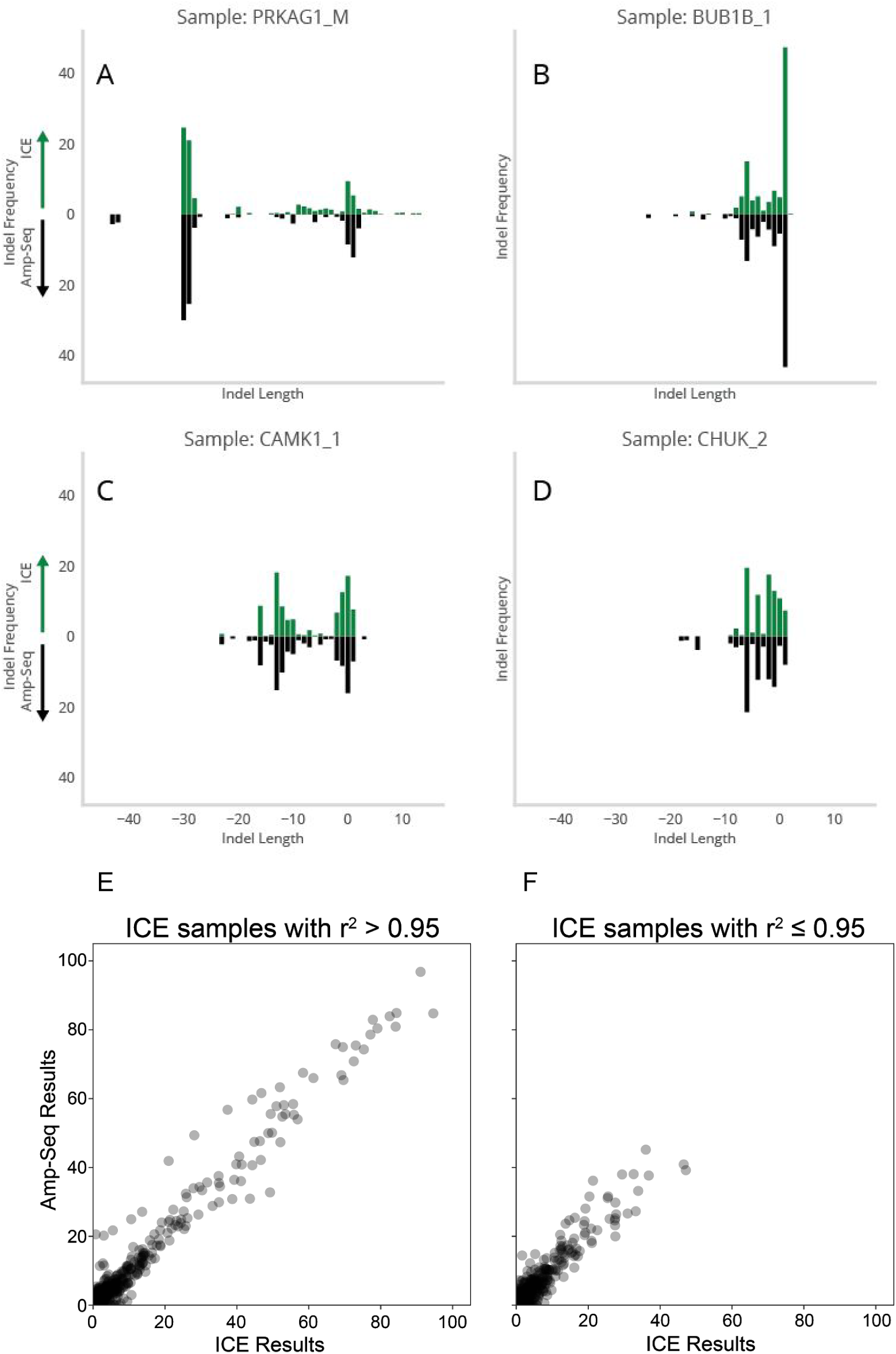
Comparison of Amp-Seq results and ICE results for 92 samples. **A-D)** the amplicon sequencing and ICE results for indel distributions for four samples. **E-F)** Comparisons of all of the pairwise points from the indel distributions. The correlation of Amp-Seq with ICE is r^2^= 0.96 for ICE samples with a high-quality analysis (Pearson r^2^ > 0.95; E). ICE results with a lower quality score (r^2^ ≤ 0.95; F) are less correlated (Pearson r^2^= 0.88), but still informative. Combining across all samples r^2^= 0.93. For each sample, the averages of up to four ICE replicates are compared to the results from the Amp-Seq run.

While specific indel frequencies correlated well between Amp-Seq and ICE, it became clear that there was little correlation in terms of the absolute number of non-zero indels (Fig 6A). This was primarily due to ICE finding indels that were not present in Amp-Seq. Our assumption with these spurious indels identified by ICE was that the regression stage was overfitting the underlying noise associated with Sanger sequencing and ascribing editing contributions to what was instrument background. We looked to address these false-positives (identified contributions that were not present in Amp-Seq) with regularization. We found that lasso regression with an L1 parameter of 0.8 performed well at decreasing the number of false positives detected by ICE, while tightening the correlation with the Amp-Seq results (Fig 6B). Beyond the absolute number of non-zero contributions, regularization resulted in an indel profile more closely resembling that of Amp-seq across tested edits (Fig 6C). Together, we found regularization to be an important addition to ICE given the underlying noise in Sanger sequencing data.

**Figure 6.**
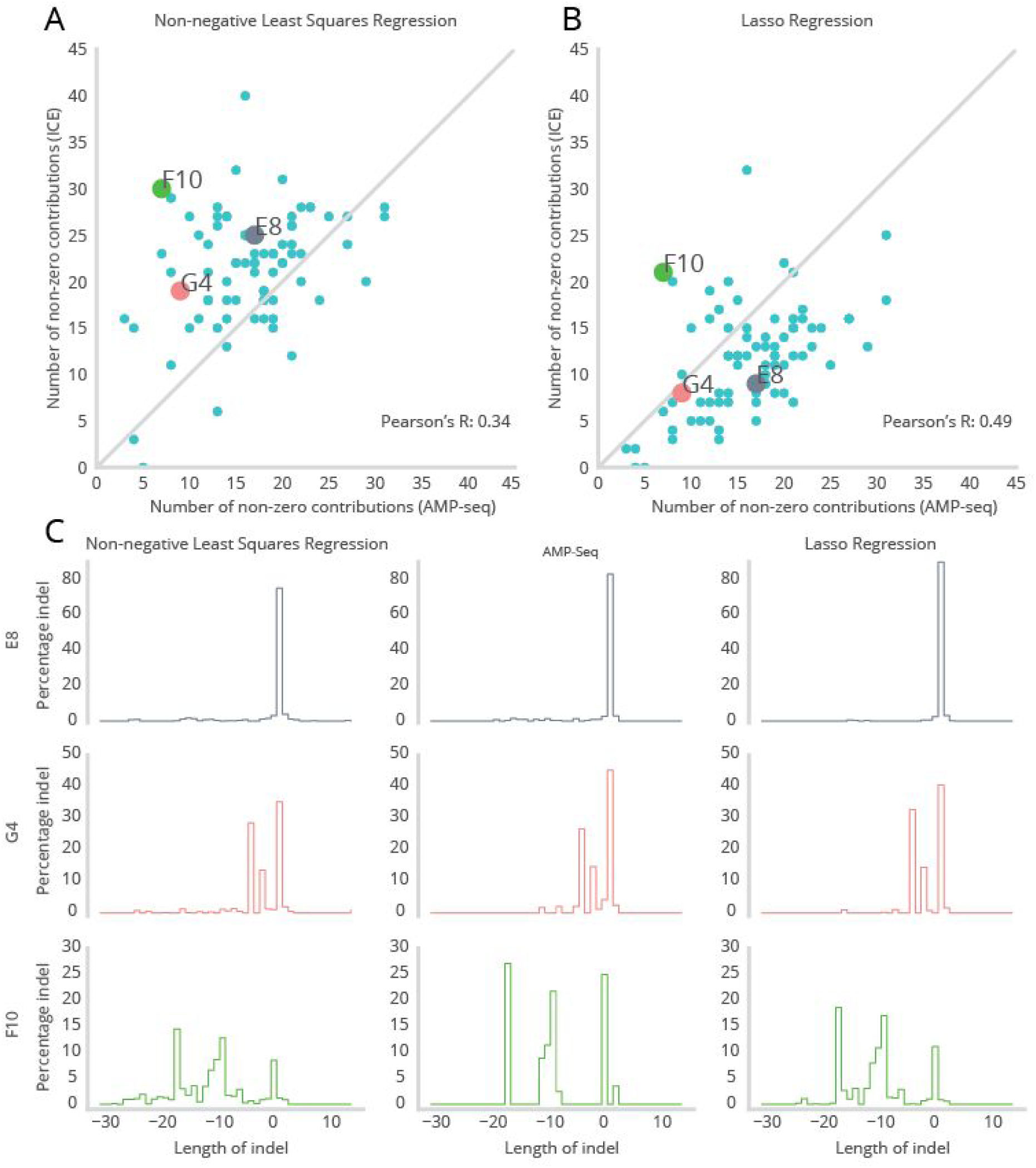
Non-Negative Least Squares and Lasso Regression when comparing ICE vs Amp-seq data. **A)** A non-negative least squares regression of the number of non-zero contributions across 79 discrete edits when sequenced with Sanger (plus ICE analysis) versus AMP-seq. The gray line represents a perfect 1:1 correlation. **B)** Sanger data collected for part A and analyzed with ICE using Lasso regression and an L1 hyperparameter value of 0.8. **C)** Indel profiles of anecdotes highlighted above showing changes in relative frequencies of particular indels in both regression algorithms compared to AMP-seq.

To specifically test ICE’s ability to accurately estimate rates of homology-directed repair (HDR), we performed amplicon sequencing on an additional 45 samples. These samples targeted 15 different cut sites and utilized donor templates with a range of insert sequence sizes (0-36 bp). We found a high correlation between Amp-Seq and ICE results for both HDR and NHEJ editing outcomes (Fig 7).

**Figure 7.**
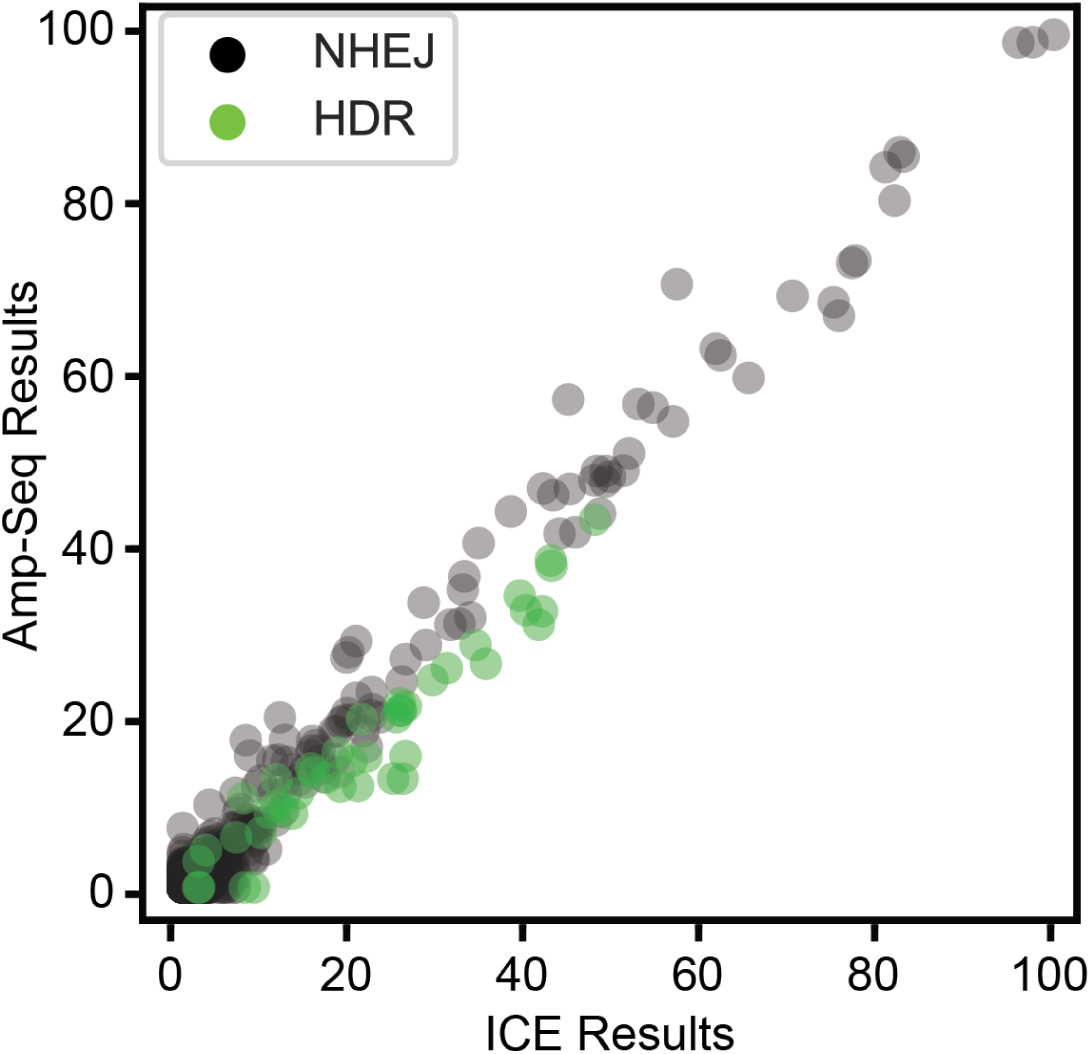
Amp-Seq results for HDR experiments were compared with ICE results for 45 samples. Each data point represents an indel for each sample (HDR= green, NHEJ= gray). The overall correlation of Amp-Seq with ICE is r^2^= 0.97.

To validate the ability of ICE to distinguish between genetic variants, we sequenced mixes of DNA that simulate a range of variant outcomes. We amplified the locus containing the SNP rs2072579 from the HEK293 cell line and George Church’s PGP1 iPSC line. Sanger sequencing confirmed the samples are homozygous and different at that position. We then quantified the amplicons with a Fragment Analyzer, mixed them in different ratios (5%,10%, 20%, 40%, 60%, 8%, 90%, 95% of PGP1 in the mixture), and sequenced the mixed samples. The sequencing data were then analyzed with ICE simulating an experiment in which the HEK293 cell line (C/C) is edited to have a homozygous G/G at SNP rs2072579. The predictions are highly correlated with the expected percentages (Fig 8).

**Figure 8.**
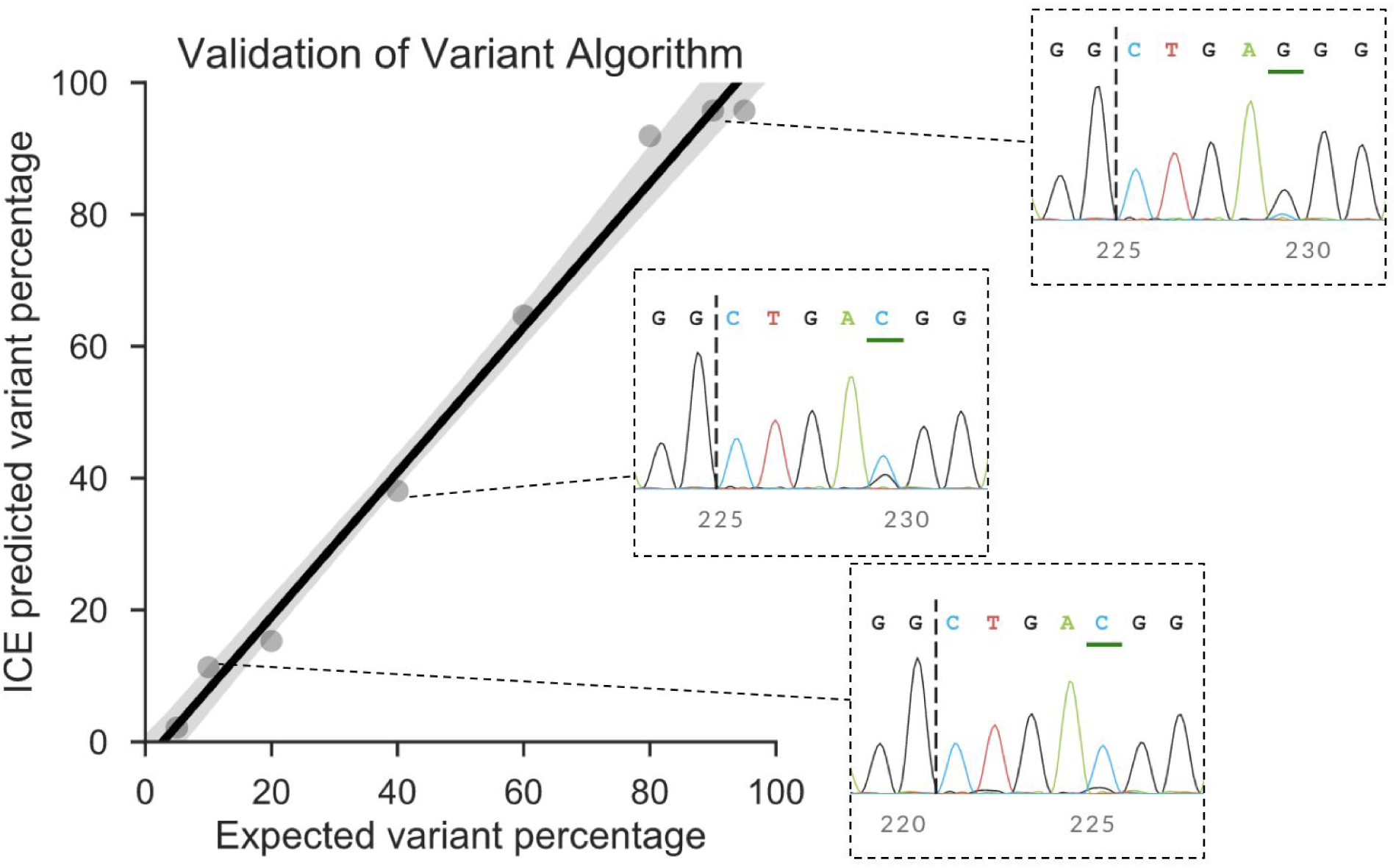
ICE variant results correlate with expected variant percentages (r^2^= 0.99).

## Discussion

Here we present our software tool, ICE, which uses Sanger sequencing data from CRISPR edited samples to quantify the identity and prevalence of edits resulting from NHEJ and HDR. ICE can be used to analyze editing outcomes from experiments that use an individual guide or multi-guide (up to 3 guides per gene) approaches. A major benefit of the ICE workflow is that it is a fast and robust assay that enables experimenters to easily optimize gene editing experiments. While Amp-Seq has better sensitivity and quantitation, Sanger sequencing still remains a more widely accessible, faster, and cheaper method. Moreover, we show a high correlation of ICE results with Amp-Seq results for 92 samples, suggesting that ICE can provide a reliable substitute in the vast majority of cases.

In comparison to TIDE, ICE is able to analyze more types of experiments, requires no subjective parameter tuning, and has comparable results. These advantages result in a webtool and software package that is able to easily process hundreds of CRISPR editing results in a reproducible manner. Additionally, ICE-D provides a form of insurance for the ICE proposing process by being able to capture unexpectedly large indels. We have validated the robustness of the ICE tool by running analyses for thousands of Sanger files in one batch.

Our approach will allow one experimental workflow to be able to analyze HDR and base editing experiments. ICE offers the benefit of not requiring laborious lab-work to construct a synthetic standard and a third Sanger sequencing, unlike TIDER [4].

There are some general assumptions made by both Sanger-based CRISPR analysis tools (ICE and TIDE) that may limit their precision. Both make the assumption that the peak signal *S* for different bases at each position is linearly proportional to the molarity of the base *m* with the relationship *S*=*bm*. Critically, the coefficient *b* is assumed to be the same for all bases. However, the peak height and phasing for a particular base in the Sanger trace is a function of the local sequence context. This could result in sequences where the molar ratios of bases present at a given position are not reflected by the Sanger signal ratios. Because base editing and HDR rely on the signal from single base positions, the peak height and phasing assumptions may have a larger adverse effect. It may be possible to better model the expected Sanger sequencing trace by using the approaches in [2]. However, the high correlation between ICE and Amp-Seq indicates that this assumption does not affect ICE’s ability to predict insertions and deletions. We suspect that because an indel affects the signal for all bases downstream, the effect of peak signal variance cancels out over many bases.

A caveat specific to the ICE multi-guide edit analysis is that the edit proposal process assumes two cuts can happen in close proximity. For example, if there are two guides with cut sites of *n* and *n*+1 in the genome, the model will generate an edit proposal where both guides cut and dropout the intervening base. However, we know this proposal is impossible as the nuclease cannot cut in parallel due to sterics and cannot cut serially as the guide sequence in the genome would have been destroyed. The addition of this constraint and other constraints that account for biological mechanisms will make it possible to bias the edit proposal process or the regression in favor of the correct sequences.

ICE offers a new and robust method for analyzing CRISPR editing experiments. ICE can detect successful edits in just a few days after transfection, as has been validated on thousands of samples. We found that ICE is able to offer results comparable to Amp-Seq, but at a significant reduction in cost and time. The ICE workflow offers several advantages over the current state-of-the-art alternatives by offering a robust and reproducible way to analyze editing experiments. It also is the only tool that can analyze multi-guide editing and requires less work to analyze HDR experiments. Because ICE reduces the labor, cost, and time associated with CRISPR experiments, analysis is no longer a limiting factor for precision genome editing.

## Appendix A: Examples of sequences predicted by ICE and those detected by Amp-Seq

Here we use bold and underlined text to match sequences in the ICE output and in the Amp-Seq contigs assembled from Amp-Seq. We also report the proportion of the sample that each approach predicts for the sequences.

### Example 1: UCK2 multi-guide editing

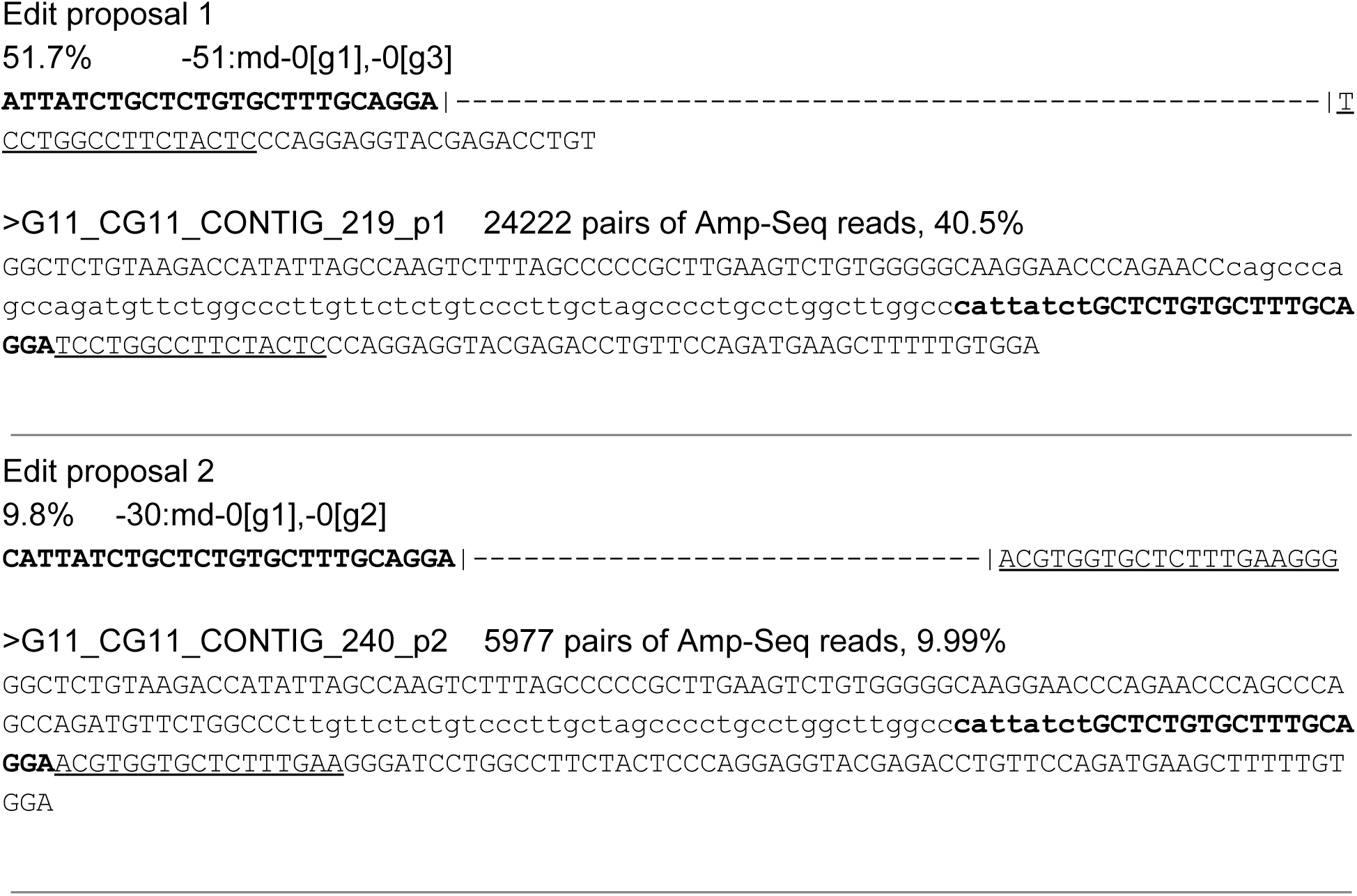

### Example 2: IRAK4 multi-guide editing

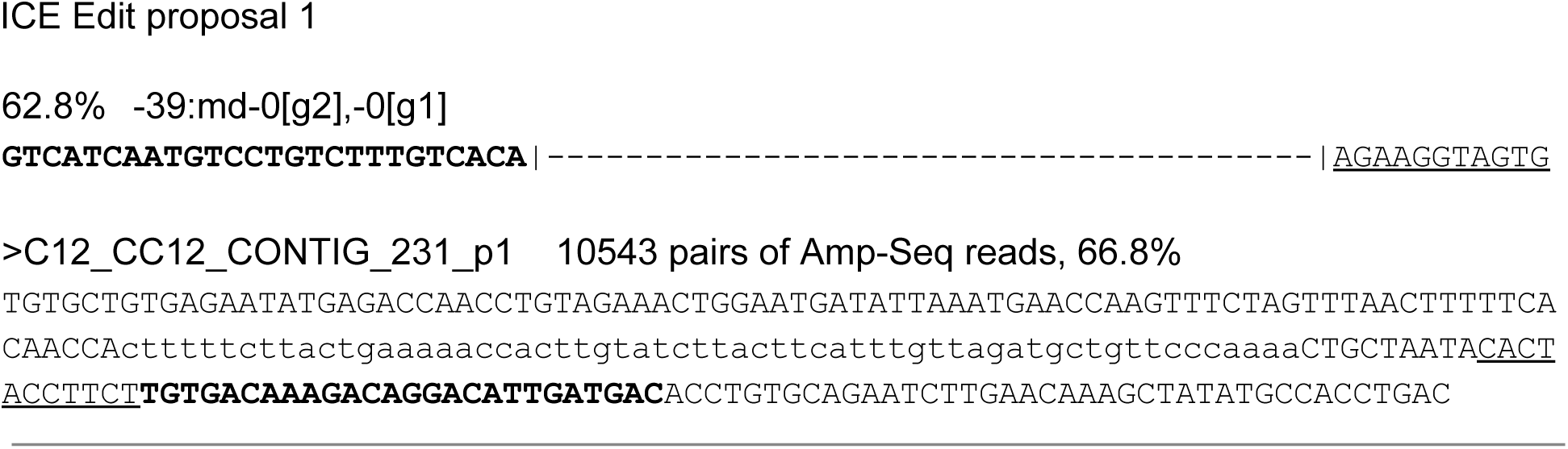

### Example 3 STK4

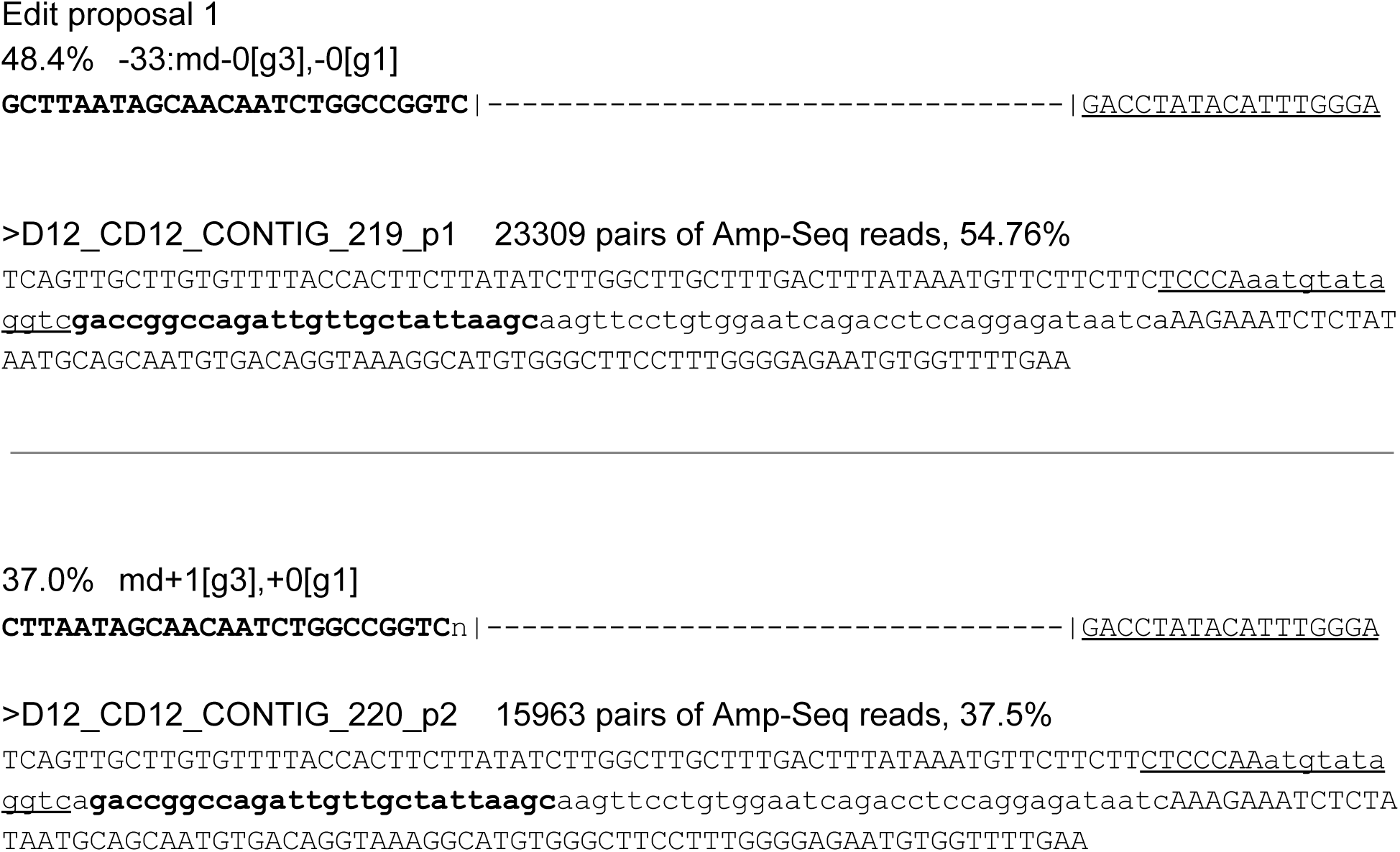

